# Photosensitized Electrospun Nanofibrous Filters for Capturing and Killing Airborne Coronaviruses under Visible Light Irradiation

**DOI:** 10.1101/2021.07.29.454404

**Authors:** Hongchen Shen, Zhe Zhou, Haihuan Wang, Mengyang Zhang, Minghao Han, Yun Shen, Danmeng Shuai

## Abstract

To address the challenge of the airborne transmission of SARS-CoV-2, photosensitized electrospun nanofibrous membranes were fabricated to effectively capture and inactivate coronavirus aerosols. With an ultrafine fiber diameter (∼ 200 nm) and a small pore size (∼ 1.5 µm), the optimized membranes caught 99.2% of the aerosols of the murine hepatitis virus A59 (MHV-A59), a coronavirus surrogate for SARS-CoV-2. In addition, rose bengal was used as the photosensitizer for the membranes because of its excellent reactivity in generating virucidal singlet oxygen, and the membranes rapidly inactivated 98.9% of MHV-A59 in virus-laden droplets only after 15 min irradiation of simulated reading light. Singlet oxygen damaged the virus genome and impaired virus binding to host cells, which elucidated the mechanism of disinfection at a molecular level. Membrane robustness was also evaluated, and no efficiency reduction for filtering MHV-A59 aerosols was observed after the membranes being exposed to both indoor light and sunlight for days. Nevertheless, sunlight exposure photobleached the membranes, reduced singlet oxygen production, and compromised the performance of disinfecting MHV-A59 in droplets. In contrast, the membranes after simulated indoor light exposure maintained their excellent disinfection performance. In summary, photosensitized electrospun nanofibrous membranes have been developed to capture and kill airborne environmental pathogens under ambient conditions, and they hold promise for broad applications as personal protective equipment and indoor air filters.

**Synopsis:** Photosensitized electrospun nanofibrous filters with excellent capture-and-kill performance against coronaviruses were designed and implemented to prevent the airborne transmission of COVID-19.

**Table of Contents:** 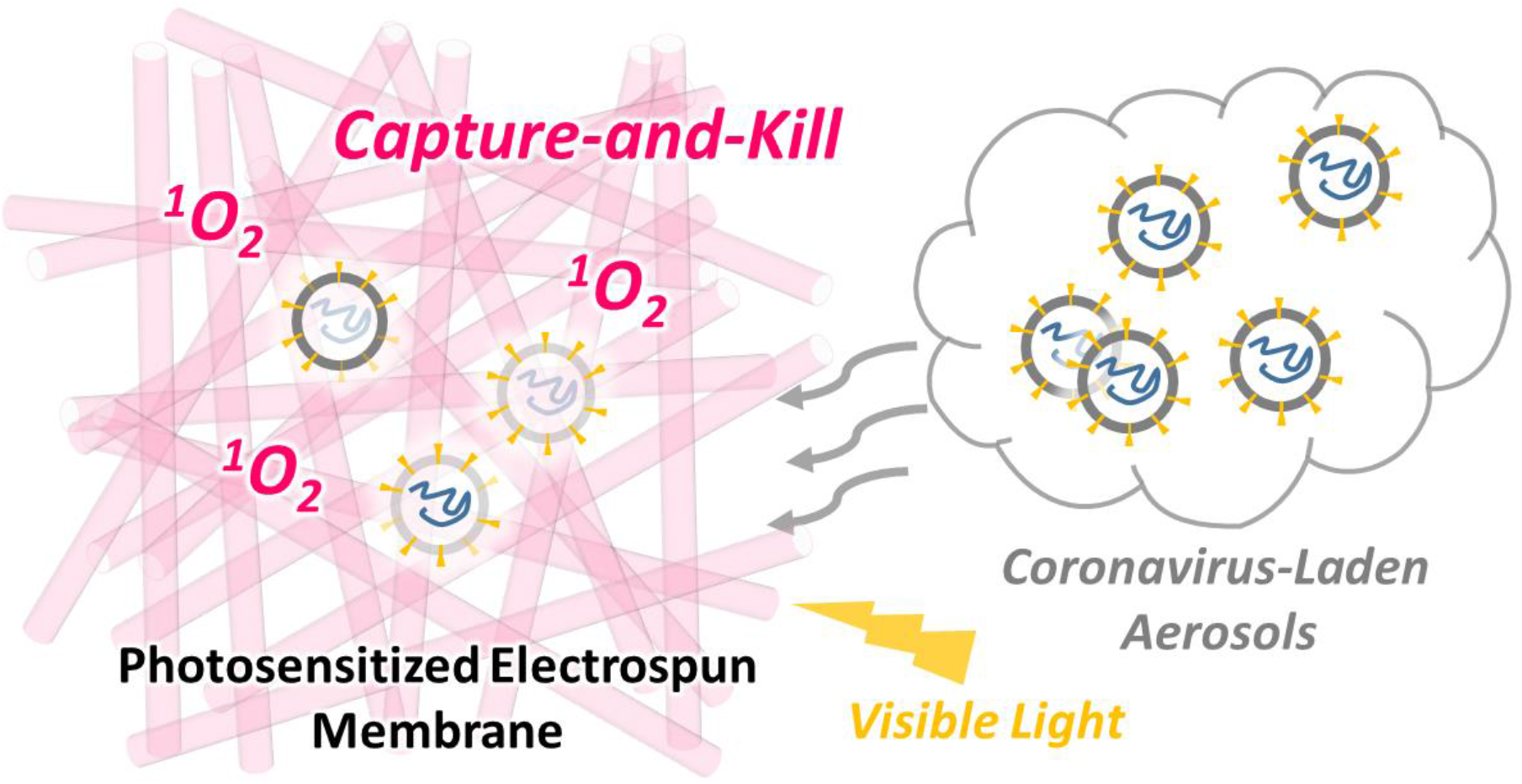

## 1. Introduction

Airborne transmission of SARS-CoV-2 has been recognized as an important route for spreading COVID-19 by the World Health Organization and the U.S. Centers for Disease Control and Prevention.^1,2^ SARS-CoV-2 aerosols could suspend, accumulate, and remain infectious in the air for a long duration up to hours.^3^ To reduce the transmission of SARS-CoV-2 through aerosols, physical barriers like face masks and indoor air filters have been successfully implemented. In particular, numerous studies have emphasized the important role of wearing masks in reducing the spread of COVID-19.^4–6^ Electrospinning has emerged as a promising nanotechnology for developing non-woven, ultrafine fibrous membranes that are excellent for removing aerosols.^7–9^ The electrospun membranes have a reduced pore size (tens of nanometers to several micrometers), an increased specific surface area, and enhanced porosity to enable efficient aerosol filtration and a low pressure drop in filtration.^7^ Furthermore, the surface and volume charges within the electrospun membranes improve aerosol filtration through electrostatic attraction.^10,11^ Particularly, our previous study has underscored that the electrospun membranes caught up to 99.9% of coronavirus aerosols whose size was within or close to that of the most penetrating particles in mechanical air filtration.^12^ However, like most masks and air filters currently used on the market, electrospun membranes only physically capture viral aerosols but they do not inactivate the viruses, which could lead to secondary contamination and potential infection (e.g., via the contact of the contaminated masks/air filters or viruses dislodged from the masks/filters).^13^

The goal of our study is to (i) develop highly efficient and robust photosensitized electrospun nanofibrous membranes that can both physically capture and chemically disinfect coronavirus aerosols, and (ii) elucidate the mechanism of photooxidation and inactivation of coronaviruses at a molecular level. We selected dye photosensitizers as antimicrobial additives for electrospinning, because they produce highly reactive singlet oxygen (^1^O_2_) for effective and rapid virus disinfection, they can be excited under the irradiation of visible light that is readily available in indoor environments, they are low cost, and some of them have been approved by the U.S. Food and Drug Administration for biomedical applications and thus pose little to no health risks to humans (e.g., rose bengal (RB)). Our membranes enable a self-cleaning feature for disinfecting coronaviruses under ambient conditions, and photosensitized disinfection does not reduce the aerosol filtration efficiency of the membranes. The photoreactive electrospun membranes can find broad air filtration applications, such as face masks, respirators, and indoor air filters, and they are developed for the first time for preventing the airborne transmission of SARS-CoV-2 and beyond.

We first optimized the composition for preparing the photosensitized electrospun nanofibrous membranes to yield the best NaCl aerosol filtration efficiency, the lowest pressure drop in filtration, and the highest reactivity for producing ^1^O_2_ under visible light irradiation. Next, we tested the filtration efficiency and inactivation kinetics of coronavirus aerosols to best understand performance of the optimum membranes for controlling the airborne transmission of COVID-19. We then conducted a thorough mechanistic investigation on how the membranes inactivated coronaviruses, damaged viral genome, and impaired viral lifecycle in host cells. Last but not least, to understand the lifetime of the membranes for practical engineering applications, we aged the membranes under continuous light exposure up to 7 days and evaluated their long-term performance of coronavirus aerosol filtration and coronavirus disinfection. Our promising results highlighted that the photosensitized electrospun nanofibrous membranes showed outstanding filtration performance for removing coronavirus aerosols, they rapidly disinfected coronaviruses under visible light irradiation, and they maintained excellent performance when used in indoor environments for a long duration.

## 2. Materials and Methods

### 2.1 Fabrication of photosensitized electrospun membranes

A series of two-layer photosensitized membranes were fabricated by electrospinning. Briefly, a homogeneous electrospinning dope solution containing *x* wt% of polyvinylidene difluoride (PVDF) (*x* = 8-20) and a dye photosensitizer in *N,N*-dimethylformamide/acetone (7/3, v/v) was electrospun onto one layer of polypropylene (PP) fabrics (VWR^®^ Basic Protection Face Mask). During the electrospinning of 10-20 wt% of PVDF, the solution feeding rate, electric field, and electrospinning duration was maintained at 0.6 mL h^-1^, 1 kV cm^-1^, and 20 min, respectively. For electrospinning 8 wt% of PVDF, the solution feeding rate, electric field, and electrospinning duration was kept at 0.4 mL h^-1^, 1 kV cm^-1^, and 30 min, respectively, to reduce bead formation. The dyes including 0.3 wt% of RB, 0.015 wt% of methylene blue hydrate (MB), 0.3 wt% of crystal violet (CV), 0.015 wt% of (-)-riboflavin (RF), and 0.003 wt% of toluidine blue O (TBO) were used. Mass percentage of PVDF and the dyes was calculated with respect to the total mass of the electrospinning dope solution, and the maximum dye concentration was selected based on dye solubility in the solution. The two-layer photosensitized electrospun membranes were denoted as PVDF*x*-dye name (e.g., PVDF15-RB). Membranes that were electrospun from 15 wt% of PVDF without dyes onto a layer of PP fabrics (VWR^®^ Basic Protection Face Mask) were also fabricated for comparison (i.e., PVDF15).

Three-layer photosensitized electrospun membranes (here referred to as sandwiched membranes), which consisted of an additional PP fabric top layer (Amazon, skin friendly non-woven fabrics) on the top of the two-layer photosensitized electrospun membranes, were particularly assembled for aerosol filtration and pressure drop tests. The sandwiched structure protected the electrospun layer from destruction during and after filtration (e.g., removal from the filter holder). Based on our previous study, the PP fabrics had negligible impact on aerosol removal and pressure drop, because of their very large pore size compared with the aerosol size (∼120 µm versus < 2 µm) and high porosity.^12^

### 2.2 Characterization of photosensitized electrospun membranes

The nanofiber diameter of electrospun membranes was characterized by scanning electron microscopy (SEM, FEI Teneo LV). For each membrane, at least 50 fibers were selected for analyzing the diameter. The pore size of two-layer photosensitized electrospun membranes was characterized by a gas liquid porometry method (POROLUXTM 100/200/500, shape factor of 0.715, APTCO Technologies LLC, Belgium). The pressure drop of sandwiched photosensitized electrospun membranes was determined with a face velocity of 5.3 cm s^-1^. Dye leaching from two-layer photosensitized electrospun membranes was estimated by optical absorbance measurements (UV-vis spectrophotometer, Hach DR6000), and details are in **Text S2**. ^1^O_2_ production of the two-layer photosensitized electrospun membranes was quantified in a liquid setup containing furfuryl alcohol (FFA, 1 mM) under the exposure of both simulated reading light (100% 7W white LED with a lamp-to-membrane distance of 15 cm) and simulated indoor light (3% 7W white LED with a lamp-to-membrane distance of 33 cm), and the steady-state ^1^O_2_ concentration ([^1^O_2_]_ss_) was calculated by dividing the measured first-order decay rate constant of FFA by the second-order reaction rate constant between ^1^O_2_ and FFA (1.2 × 10^8^ M^-1^ s^-1^). Simulated reading light had a higher photon flux and optical power density than those of simulated indoor light, and they were recorded in **Figure S1** and **Table S1** (AvaSpec-2048 Fiber Optic Spectrometer). FFA was monitored by high performance liquid chromatography (Shimadzu LC-20AT Prominence).

### 2.3 Determination of filtration efficiency for removing NaCl and coronavirus aerosols

NaCl solution (0.1 M) and murine hepatitis virus A59 (MHV-A59) in water (∼10^6^-10^7^ gene copies mL^-1^, diluted from the virus stock by ∼50 times with nuclease-free water) were used for aerosolization and filtration tests. MHV-A59 was selected because it is a β-coronavirus that shares the same family and the size with SARS-CoV-2 (85 nm of MHV-A59 versus 50-200 nm of SARS-CoV-2).^14^ Details of MHV-A59 propagation are in **Text S3**. Only for the aerosol size characterization, aerosols were generated from ultrapure water containing polystyrene and silica nanoparticles (00876-15, Polysciences; SISN100-25M, nanoComposix; both were 100 nm) to simulate MHV-A59 aerosols, because both nanoparticles and the MHV-A59 had a similar size and concentration during aerosolization and the two different nanoparticles had representative hydrophobicity/hydrophilicity. In the filtration tests (i.e., filter-on), a portion of NaCl or MHV-A59 aerosols were captured by the sandwiched membranes, whereas the penetrating aerosols were retained by an impinger. In the control experiments (i.e., filter-off), no filter was in place and all generated aerosols were retained by the impinger. The filtration efficiency was determined based on the difference of the amount of NaCl or MHV-A59 in the impinger between filter-off and filter-on experiments over the amount of NaCl or MHV-A59 in the impinger in the filter-off experiment. At least duplicates were conducted for each filtration test and control experiment. Details of the setup and experimental conditions are described in our previous study and also in **Text S4**.^12^ NaCl was quantified by ion chromatography (Dionex ICS-1100), and the amount and infectivity of MHV-A59 were quantified by the reverse transcription-quantitative polymerase chain reaction (RT-qPCR) and the integrated cell culture-reverse transcription-quantitative polymerase chain reaction (ICC-RT-qPCR), respectively.

### 2.4 RT-qPCR quantification of coronaviruses

MHV-A59 collected in the impinger was concentrated by centrifugal ultrafiltration (Nanosep, 300 kDa, Pall Laboratory), and next proceeded for RNA extraction by a Zymo Quick-RNA Viral Kit (R1035). MHV-A59 was quantified by RT-qPCR by amplifying a fraction of its ORF5 gene for the structural protein M.^15,16^ The TaqMan™ Fast Virus 1-Step Master Mix Kit (Thermo Fisher Scientific Inc., 4444432) was used for RT-qPCR. Details of RNA extraction efficiency, the sequence and concentration of primers, probe, and cDNA standard, reverse transcription and PCR programs, PCR amplification efficiency, positive and negative controls, and inhibition tests are summarized in **Text S5**. All the RT-qPCR data were reported following MIQE guidelines.^12,17^

### 2.5 Evaluation of coronavirus infectivity after photosensitization

Compared with conventional infectivity assays based on plaque forming units or median tissue culture infectious dose (TCID_50_), ICC-RT-qPCR is a rapid, sensitive, and reliable method for quantifying the infectivity of coronaviruses, based on RT-qPCR of viral gene copy numbers after virus replication in the host cells. The experimental conditions of ICC-RT-qPCR were optimized, and the method is valid for quantifying the infectivity of MHV-A59 because a linear correlation was observed between virus infectivity quantified by ICC-RT-qPCR and the viral load before infection determined by RT-qPCR (**Figure S2**). Special attention should also be paid for interpreting the virus infectivity quantified by ICC-RT-qPCR, because ICC-RT-qPCR compared with a conventional cell culture assay could overestimate the dosage of reactive oxygen species (ROS) to achieve the same level of virus inactivation.^18^

The infectivity of MHV-A59 captured on the photosensitized electrospun membrane was evaluated by ICC-RT-qPCR after the exposure to simulated reading light. Briefly, the viruses on the membrane were eluted in nuclease-free water, concentrated by centrifugal ultrafiltration (Nanosep, 300 kDa, Pall Laboratory), and proceeded for cell infection and RT-qPCR quantification. Details are included in **Text S6**. Unfortunately, the infectivity of MHV-A59 that was eluted from the membrane after aerosol filtration was not able to be quantified by ICC-RT-qPCR, because aerosolization lost most of the viruses (i.e., only ∼1 out of a million viruses was able to be aerosolized in our system) and a low multiplicity of infection did not allow virus propagation in the host cells.^19^ To facilitate quantifying the photoreactivity of photosensitized electrospun membranes for inactivating MHV-A59, we designed a liquid setup that virus-laden droplets were loaded on the membrane surface under the exposure of both simulated reading light and simulated indoor light (details in **Text S7**). First-order infectivity decay rate constants, *k*_*infectivity*_, were obtained from the negative slope of the linear regression of the natural logarithm of MHV-A59 infectivity versus the light exposure duration. The liquid setup was amended with a high viral load, and it provided reliable and quantifiable infectivity of the virus after light exposure. The liquid setup could best simulate the scenario of coronavirus inactivation when virus-laden respiratory droplets are captured on the masks. Control experiments that evaluated MHV-A59 inactivation by PVDF15 under the irradiation of simulated reading light and by PVDF15-RB in the dark were also conducted. Triplicates were conducted for MHV-A59 inactivation by PVDF15-RB under light exposure, and duplicates were conducted for the control experiments.

### 2.6 Evaluation of coronavirus gene damage after photosensitization

After photosensitization, the ORF5 gene damage of both filter-captured MHV-A59 aerosols and MHV-A59 in droplets was evaluated by RT-qPCR. The same experimental setup for investigating MHV-A59 infectivity was used, as described in **Section 2.5**, and MHV-A59 aerosols captured on the sandwiched PVDF15-RB and MHV-A59 droplets on PVDF15-RB were subjected to the irradiation of simulated reading light. The samples were next collected at different time intervals for RT-qPCR quantification, as described in **Section 2.4**. Photooxidation damaged the viral genes and prevented its RT-qPCR quantification, and only the intact genes were able to be determined. Control experiments that evaluated the ORF5 gene damage of captured MHV-A59 aerosols on PVDF15 under the irradiation of simulated reading light and on PVDF15-RB in the dark were also conducted. First-order ORF5 gene damage rate constants, *k*_*gene*_, were obtained from the negative slope of the linear regression of the natural logarithm of the ORF5 gene copy number quantified by RT-qPCR versus the light exposure duration. Triplicates were conducted for the ORF5 gene damage of MHV-A59 by PVDF15-RB under light exposure, including both viral aerosols and droplets; and duplicates were conducted for the control experiments.

### 2.7 Evaluation of coronavirus’ lifecycle after photosensitization

The impact of photosensitization on coronavirus’ lifecycle in host cells, including virus binding and internalization, was investigated. The same liquid setup as described in **Section 2.5** was used, and MHV-A59 droplets on PVDF15-RB were subjected to the irradiation of simulated reading light (details in **Text S8**). Incubation MHV-A59 with L-929 cells at 4 °C only allowed virus binding, whereas subsequent increase of the incubation temperature to 37 °C permitted virus internalization.^20–22^ First-order rate constants of apparent damage to coronavirus’ lifecycle were obtained from negative slope of the linear regression of the natural logarithm of RT-qPCR quantified viruses bound to and internalized into the cells versus the light exposure duration, and they are denoted as *k*_*app_binding*_ and *k*_*app_internalization*_, respectively. The rate constants of true damage to virus binding and internalization were calculated as *k*_*binding*_=*k*_*app_binding*_-*k*_*gene*_ and *k*_*internalization*_=*k*_*app_internalization*_-*k*_*app_binding*_, respectively.^18^ The correction for *k*_*app_binding*_ was to distinguish the decrease of ORF5 gene PCR signal due to ORF5 gene damage by photooxidation and the loss of virus binding function by the same treatment; and the correction for *k*_*app_internalization*_ was to exclude the contribution of the decrease of ORF5 gene PCR signal due to damaged virus binding by photosensitization. Triplicates were conducted for the damage of coronavirus’ lifecycle after photosensitization.

### 2.8 Aging and robustness of photosensitized electrospun membranes

Fresh two-layer photosensitized electrospun membranes were aged under different light exposure to explore their robustness and lifetime for capturing and killing coronaviruses. Briefly, PVDF15-RB was exposed to indoor light (fluorescent light in the laboratory) and simulated indoor light up to 7 days and outdoor sunlight up to 4 days, and the aged membranes were denoted as PVDF-RB-I, PVDF-RB-S, and PVDF-RB-O, respectively. The light spectrum and intensity for aging are included in the **Figure S1** and **Table S1**. Particularly, indoor light and simulated indoor light had almost identical photon flux and optical power density. After aging, the membranes were characterized by SEM for morphology and nanofiber diameter, and their production of ^1^O_2_ under the irradiation of both simulated reading light and simulated indoor light was also measured, as described in **Section 2.2**. Aged membranes were also used for testing the filtration efficiency for removing MHV-A59 aerosols and the inactivation of MHV-A59 droplets, as described in **Sections 2.3** and **2.5**.

### 2.9 Data analysis

Student’s t test was utilized to determine whether the calculated first-order reaction rate constants of infectivity decay, ORF5 gene damage, and the damage of virus binding and internalization were different from 0. Student’s t test was also used for the statistical comparison of the difference between two first-order reaction rate constants, [^1^O_2_]_ss_, and aerosol filtration efficiencies. All *p* values < 0.05 were considered statistically significant.

## 3. Results and Discussion

### 3.1 Optimizing photosensitized electrospun membranes

All photosensitized electrospun membranes showed the color of the photosensitizers, in contrast to the white bare PVDF membranes, indicating the successful incorporation of photosensitizers into the PVDF matrix (**Figure 1a**). The loading of photosensitizers was maximized based on their solubility in the electrospinning dope solution to ensure the best photoreactivity for inactivating coronaviruses. The hydrophilic photosensitizers were embedded in the hydrophobic PVDF, which minimized photosensitizer leaching. Specifically, only 2.20 wt% of RB was released when PVDF15-RB was immersed in water for 6 h, suggesting that the moisture from human breath or ambient air would not induce significant photosensitizer leaching when the membranes are used as masks and indoor air filters. To select the photosensitizer with the best photoreactivity, [^1^O_2_]_ss_ produced from the electrospun membranes loaded with different photosensitizers was characterized. Under the exposure of simulated reading light with a higher light intensity, PVDF15-RB produced a [^1^O_2_]_ss_ of (6.94 ± 2.19) × 10^−13^ M, which was 2.31, 3.81, 8.68, and 4.08 times higher than that of PVDF15-MB, PVDF15-CV, PVDF15-RF, and PVDF15-TBO (*p* < 0.05, **Figure S4**). Under the irradiation of simulated indoor light with a lower light intensity, PVDF15-RB and PVDF15-MB outperformed other photosensitized electrospun membranes for producing ^1^O_2_ (*p* < 0.05, **Figure S4**). Therefore, RB was identified as the most photoreactive dye additive for membrane fabrication. The fact that PVDF15-RB has the highest photoreactivity could be attributed to the increased photon absorption under white LED light irradiation (maximum absorption wavelength of RB is 546 nm) and a high quantum yield of the intersystem crossing of RB for producing ^1^O_2_.^23,24^

**Figure 1.**
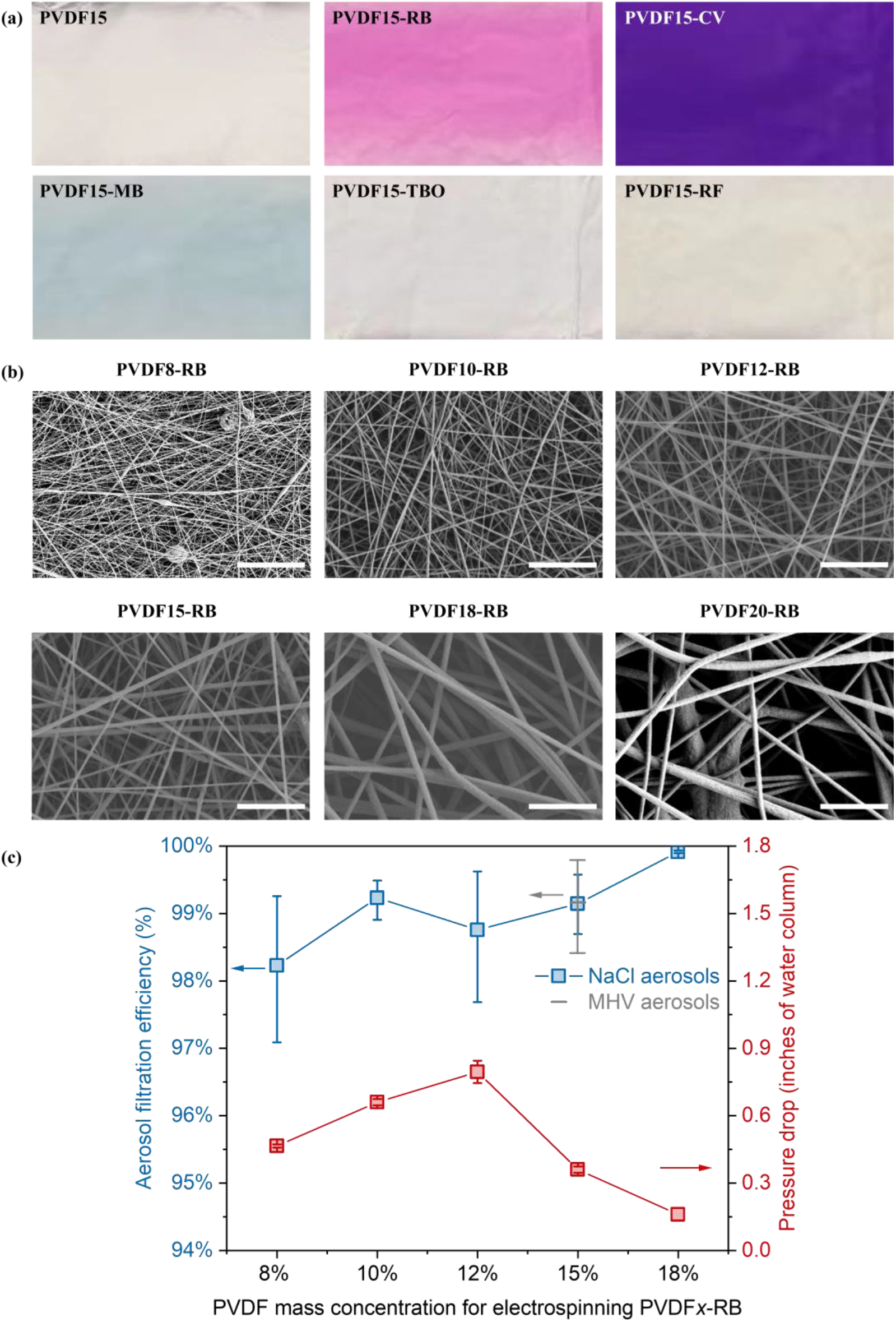
(a) Photosensitized nanofibrous membranes electrospun with various photosensitizers; (b) SEM for the RB-sensitized electrospun membranes with different PVDF concentrations. Scale bars in SEM images are 5 µm ; (c) NaCl and MHV-A59 aerosol filtration efficiency and pressure drop in filtration for RB-sensitized electrospun membranes with different PVDF concentrations (PVDF*x*-RB). Blue squares and error bars in the filtration efficiency graph represent average and maximum/minimum values of duplicates, and red squares and error bars in the pressure drop graph represent the average and standard deviation of triplicates.

The PVDF concentration for developing photosensitized electrospun membranes was next optimized to improve filtration performance. As the key factors determining the aerosol filtration efficiency, the nanofiber diameter and pore size of the membranes were characterized at different polymer concentrations. The nanofiber diameter of PVDF8-RB, PVDF10-RB, PVDF12-RB, PVDF15-RB, PVDF18-RB, and PVDF20-RB was 60 ± 16, 116 ± 24, 164 ± 48, 196 ± 44, 521 ± 191, and 423 ± 114 nm, respectively (**Figure 1b**). Generally, the nanofiber diameter increased with the increase of the PVDF concentration from 8 to 15 wt% and the membranes possessed uniform nanofibers. When PVDF concentration reached to 18 and 20 wt% for electrospinning, the fibers were not uniform anymore and some very large fibers were present. This could be attributed to a significantly increased viscosity of the electrospinning dope solution and instable and non-continuous electrospinning jets.^25^ In addition, the nanofiber diameter and pore size of PVDF15-RB (196 ± 44 nm, 1.46 ± 0.15 µm) was smaller than those of the bare electrospun membrane of PVDF15 (315 ± 73 nm, 2.70 ± 0.20 µm), likely due to the presence of anionic RB and increased conductivity of the dope solution for electrospinning.^12,26,27^

The filtration efficiency of NaCl aerosols was further examined for those PVDF*x*-RB membranes. Our previous study has demonstrated the filtration efficiency of NaCl aerosols was a conservative indicator for understanding the removal of coronavirus aerosols.^12^ The average filtration efficiency for NaCl aerosols was 98.2%, 99.2%, 98.8%, 99.1%, and 99.9% for PVDF8-RB, PVDF10-RB, PVDF12-RB, PVDF15-RB, and PVDF18-RB, respectively (**Figure 1c**). PVDF20-RB was not tested for filtering aerosols because the membrane was apparently non-uniform after electrospinning. Pressure drop in filtration was also determined to understand the breathability or energy consumption in filtration. With the increase of the PVDF concentration from 8 to 12 wt% in the RB-sensitized electrospun membranes, the pressure drop increased from 0.47 ± 0.01 to 0.80 ± 0.05 inches of water column (inches wc). However, further increase of the PVDF concentration from 12 to 18 wt% reduced the pressure drop to 0.16 ± 0.00 inches wc (**Figure 1c**). The low pressure drop at a low PVDF concentration could be attributed to the slip effect of air molecules on the ultrafine nanofibers, whereas the low pressure drop at a high PVDF concentration could be resulted from a high membrane porosity.^28^ PVDF15-RB had the best filtration performance including a high aerosol filtration efficiency and a low pressure drop, the best photoreactivity in terms of ^1^O_2_ production, and uniform nanofibers across the whole membrane that allows manufacturing at scale. Therefore, PVDF15-RB was selected for the following study of coronavirus filtration and inactivation.

### 3.2 Photosensitized electrospun membranes for capturing and inactivating coronavirus aerosols

The optimized electrospun membrane of PVDF15-RB was further challenged by MHV-A59 aerosols, and it removed 99.2% of the viral aerosols on average (**Figure 1c**). The filtration efficiency for MHV-A59 aerosols was on par with that for removing NaCl aerosols (99.1%, *p* > 0.05), and it was much higher compared with filtration efficiency of MHV-A59 aerosols by commercial masks, including a surgical mask (98.2%), a cotton mask (73.3%), and a neck gaiter (44.9 %).^12^ The excellent aerosol filtration efficiency was resulted from the ultrafine nanofibers and the small pore size of the electrospun membrane. Our previous study also reported that PVDF15 captured 99.1% of the MHV-A59 aerosols,^12^ which was comparable with that of PVDF15-RB (*p* > 0.05). Size distribution of simulated MHV-A59 aerosols, generated from polystyrene and silica nanoparticles in water, underscored that the most dominant aerosols were in the size of 420-450 nm (**Figure S5**). Aerosol size characterization also highlighted that 79.9-89.5% of the simulated MHV-A59 aerosols were between 200 and 500 nm (**Figure S5**) and they were considered as the most penetrating aerosols in mechanical filtration.^29,30^ The optimized photosensitized electrospun membrane holds promise for effectively removing the most challenging coronavirus aerosols, which could minimize and prevent the airborne transmission of pathogens.

Beyond physical capture of the coronavirus aerosols, photosensitized electrospun membranes also inactivated the captured coronaviruses under visible light irradiation through the rapid and potent oxidation by ^1^O_2_. The captured MHV-A59 aerosols on PVDF15-RB were exposed to the irradiation of simulated reading light for 1 h, and the ORF5 gene copy number of MHV-A59 was reduced by 93.7%, giving *k*_*gene*_ of 0.0461 ± 0.0044 min^-1^ (**Figures 2a** and **2b**). In contrast, in the control experiments, no apparent ORF5 gene damage was observed when the photosensitizer or the visible light was not present (**Figure S6**). These results indicated that the photosensitized electrospun membranes damaged the coronavirus genome and could potentially inactivate the viruses upon visible light exposure. However, only a small fraction of MHV-A59 genome was quantified (∼100 bases) in the RT-qPCR, and it could not represent or reveal the damage in other regions in the whole genome (∼31.5 kb). Different regions across the viral genome can contain diverse nucleotide sequences and form unique secondary structures, which may have distinct susceptibility to ROS oxidation.^31^ A genome-wide approach that quantifies a large fraction of the genome damage by PCR could be used to reasonably predict the whole genome damage upon oxidation in the future.^32^ More importantly, the quantity of the ORF5 gene determined by RT-qPCR did not represent the viability of MHV-A59. During photosensitization, RB produces ^1^O_2_ as the key ROS for inactivating viruses, through the oxidation of genomes, proteins, lipids, and any other functional biomolecules of the viruses.^33,34^ Because ROS are broad-spectrum oxidants that damage multiple viral biomolecules at the same time and any critical damage could compromise viral viability, virus infectivity was observed to decrease more quickly compared with intact genes.^18^ Therefore, the RT-qPCR quantification of intact genes can serve as a conservative approach to understand virus infectivity, but the infectivity assay is needed to elucidate the inactivation efficiency by the photosensitized electrospun membranes.

**Figure 2.**
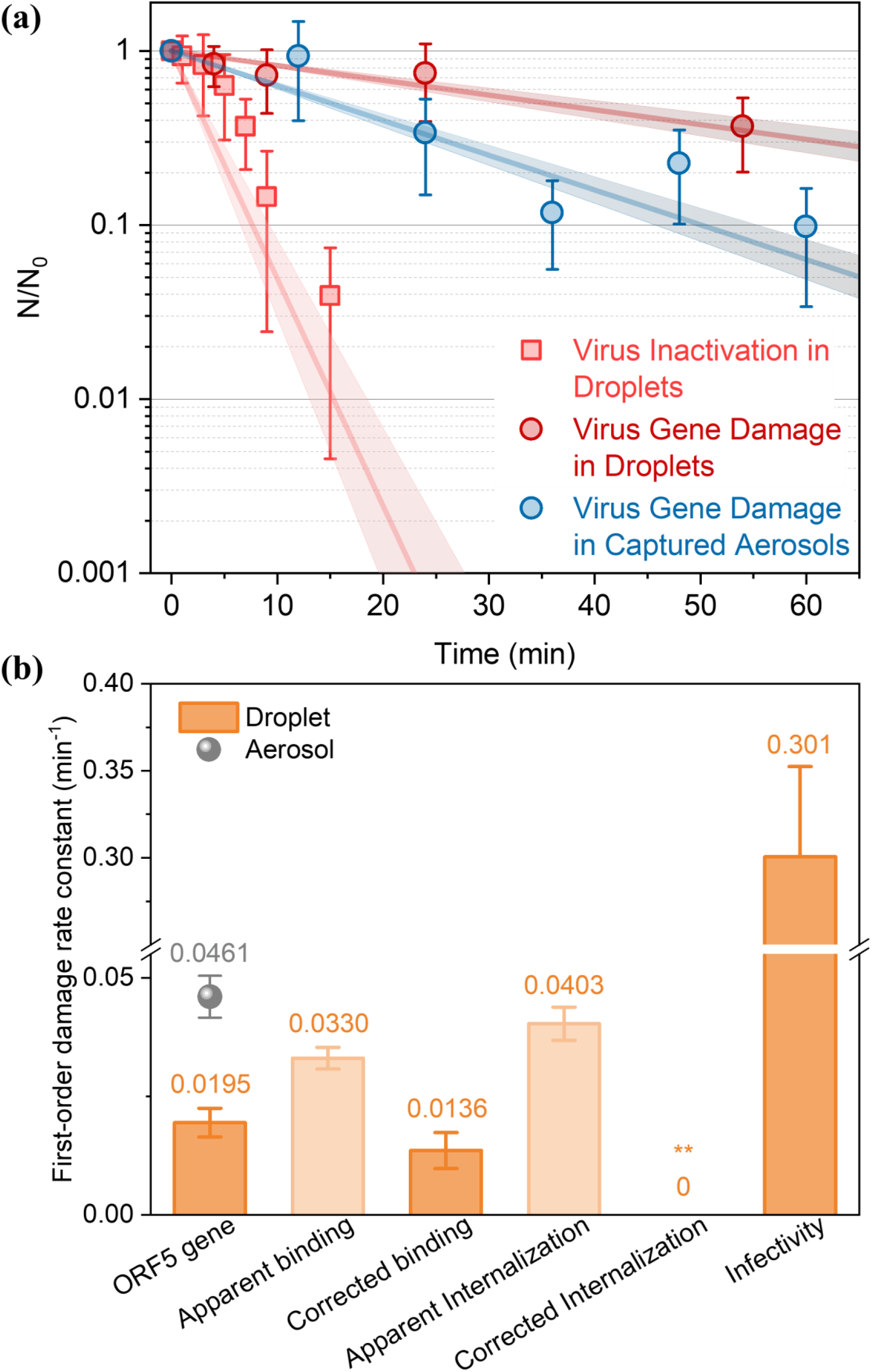
(a) First-order reaction kinetics of MHV-A59 infectivity decay in droplets and first-order reaction kinetics of MHV-A59 ORF5 gene damage in both droplets and membrane-captured aerosols. N/N_0_ represents ORF5 gene copy numbers quantified by ICC-RT-qPCR (for infectivity) or RT-qPCR (for gene damage) at light exposure duration *t* to that at light exposure duration zero. Linear regression of common logarithm of N/N_0_ versus light exposure duration and its standard errors are illustrated. Error bars represent the standard deviation of triplicates. All tests were conducted under simulated reading light exposure. (b) First-order damage rate constants for the MHV-A59 ORF5 gene, and virus binding, internalization, and infectivity under simulated reading light exposure. (**) denotes the rate constant was assigned a value of zero because there was no statistical difference between the apparent damage rate constants of internalization and binding (*p* > 0.05). Error bars represent the standard error of the negative slope of linear regression.

Though the ORF5 gene copy number of MHV-A59 aerosols was successfully quantified by RT-qPCR with a high sensitivity, virus infectivity was below the detection limit of ICC-RT-qPCR because only few coronaviruses were harvested from the membranes. The majority of coronaviruses were lost during aerosolization, since the partitioning coefficient of the coronavirus (the virus concentration in aerosols to that in the liquid solution for aerosolization) was only 5.80 ± 4.26 × 10^−7^. To overcome the challenges in quantifying the viability of coronavirus aerosols, we loaded MHV-A59 droplets (∼ 1 mm thick) on PVDF15-RB and determined virus infectivity during photosensitized disinfection. The setup best mimicked virus-laden respiratory droplets captured by the masks, and it allowed the accurate quantification of virus infectivity because of a higher concentration of MHV-A59 in the droplets than that in the aerosols. Under simulated reading light exposure, we observed significant coronavirus inactivation, with *k*_*infectivity*_ of 0.301 ± 0.052 min^-1^ (**Figure 2** and **2b**). That is said, 98.9% of coronaviruses were inactivated after only 15 min irradiation of the simulated reading light. In contrast, we did not see any noticeable coronavirus inactivation under simulated indoor light exposure up to 30 min. Control experiments of PVDF15 under simulated reading light exposure and PVDF15-RB in the dark did not inactivate the coronaviruses either (**Figure S7**). These results indicated that photosensitization with the presence of both the dye and photons is needed for coronavirus inactivation. In addition, PVDF15-RB produced more ^1^O_2_ under the irradiation of simulated reading light than simulated indoor light ([^1^O_2_]_ss_ = (6.94 ± 2.19) × 10^−13^ versus (4.95 ± 0.80) × 10^−14^ M) (**Figure S4**), and thus much more rapid coronavirus inactivation was observed under light exposure with a stronger intensity. We also found that the ORF5 gene damage rate of MHV-A59 in the droplets was lower than that of aerosolized MHV-A59 captured on the membranes after exposure to simulated reading light (*k*_*gene*_ = 0.0195 ± 0.0030 versus 0. 0461 ± 0.0044 min^-1^, *p* < 0.05, **Figure 2** and **2b**). The slower ORF5 gene damage could be attributed to the shorter lifetime and diffusion length of ^1^O_2_ in water compared with that in the air (2 μs in distilled water versus 2.80 s in the air), due to energy dissipation resulted from the collision between ^1^O_2_ and water molecules.^35,36^ The dissolved oxygen concentration in water was also lower than the oxygen concentration in the air (< 10 mg L^-1^ in the water versus 275 mg L^-1^ in the air at 25 °C), which might limit ^1^O_2_ production in the aqueous phase.^37^ Since ^1^O_2_ was the key ROS in photosensitization and it did not bias damaging the viral genome and inactivating the viruses, the liquid setup is considered as a conservative system to evaluate the inactivation of MHV-A59 aerosols captured on the photosensitized electrospun membrane. Therefore, it is reasonable to speculate that the inactivation of infectious or viable MHV-A59 in the aerosols captured on the membrane will be much faster than that in the droplets, though the infectivity of viral aerosols was not quantifiable in our study.

### 3.3 Impact of photooxidation on the lifecycle of coronaviruses

We next investigated the effect of photosensitization on the lifecycle of coronaviruses in host cells, particularly virus binding and internalization, to understand the impact of ROS on coronavirus inactivation at a molecular level. For a viable MHV-A59, the viral spike (S) protein first binds to the receptor of the host cells, murine carcinoembryonic antigen-related cell adhesion molecule 1a (mCEACAM1a), and initiates the viral lifecycle.^38^ Next, MHV-A59 transports RNA genomes into the host cells by direct fusion of the viral envelope membrane with the cell plasma membrane, or by endocytosis with subsequent fusion of the viral envelop membrane and the endosomal membrane.^39^ After internalization, viral genomes start replication as soon as sufficient machinery proteins are produced. Once progeny viruses are constructed and assembled, they are released from the old host cells and start their new lifecycle by infecting more host cells. It is clear that virus binding and internalization are key steps that determine the fate and infectivity of coronaviruses, and any damage to these biological processes could compromise the infectivity of the viruses.

We evaluated MHV-A59 binding and internalization in the liquid setup. Coronavirus binding to the L-929 cells was impaired after photooxidation, and *k*_*app_binding*_ was estimated as 0.0330 ± 0.0023 min^-1^. By excluding the contribution of ORF5 gene damage by oxidation (*k*_*gene*_ = 0.0195 ± 0.0030 min^-1^), the true decay rate constant of coronavirus binding (*k*_*binding*_) was 0.0136 ± 0.0038 min^-1^ (*p* < 0.05, **Figure 2b**). Coronavirus internalization into the L-929 cells is the subsequent step after virus binding, therefore the difference between *k*_*app_internalization*_ and *k*_*app_binding*_ is considered as the true damage to coronavirus internalization (*k*_*internalization*_). However, photooxidation did not compromise coronavirus internalization, because *k*_*app_internalization*_ and *k*_*app_binding*_ were statistically the same (0.0403 ± 0.00 35 versus 0.0330 ± 0.0023 min^-1^, *p* > 0.05, **Figure 2b**). The S protein of MHV-A59 plays important roles in both virus binding and internalization: the receptor-binding domain in subunit 1 governs virus attachment to the host cells and the fusion peptide in subunit 2 is responsible for virus internalization.^38,40 1^O_2_ produced in photosensitization non-selectively oxidizes and damages viral biomolecules, including the S protein. However, we speculate that the receptor-binding domain might be much more susceptible to ^1^O_2_ oxidation compared with the fusion peptide, because the latter contains a large amount of Ala and Gly that react with ^1^O_2_ slowly.^41,42^ It could explain why virus binding but not internalization was significantly impaired after photosensitization, and further research is required.

### 3.4 Aging of photosensitized electrospun membranes and their long-term performance

Photosensitizers are generally sensitive to long-term light exposure, because the generated ^1^O_2_ also oxidizes the photosensitizers, leads to photobleaching, and reduces the photoreactivity for the continuous production of ^1^O_2_ and virus inactivation. After 4 days of sunlight exposure (including both daytime and nighttime), the color of PVDF15-RB-O was significantly faded in comparison to fresh PVDF15-RB. Nevertheless, after 7 days of continuous irradiation of simulated indoor light and real indoor light, no apparent fading was observed for PVDF15-RB-S and PVDF15-RB-I (**Figure 3a**). ^1^O_2_ production was also characterized for all these aged membranes, and [^1^O_2_]_ss_ was (2.26 ± 0.11) × 10^−13^, (5.64 ± 1.18) × 10^−13^, and (7.75 ± 1.70) × 10^−13^ M for PVDF15-RB-O, PVDF15-RB-S, PVDF15-RB-I, respectively, under the scenario of simulated reading light exposure (**Figure S4**). Significant less ^1^O_2_ production was observed for PVDF15-RB-O when compared to PVDF15-RB (*p* < 0.05), but a similar amount of ^1^O_2_ was generated from PVDF15-RB, PVDF15-RB-S, and PVDF15-RB-I (all *p* > 0.05). Membrane morphology was also characterized, but aging under light exposure for extended duration did not change the nanofiber diameter: the diameters were 203 ± 51, 200 ± 43, and 183 ± 45 nm for PVDF15-RB-I, PVDF15-RB-S, and PVDF15-RB-O, respectively, in comparison with the fresh PVDF15-RB with a fiber diameter of 196 ± 44 nm (**Figure 3a**).

**Figure 3.**
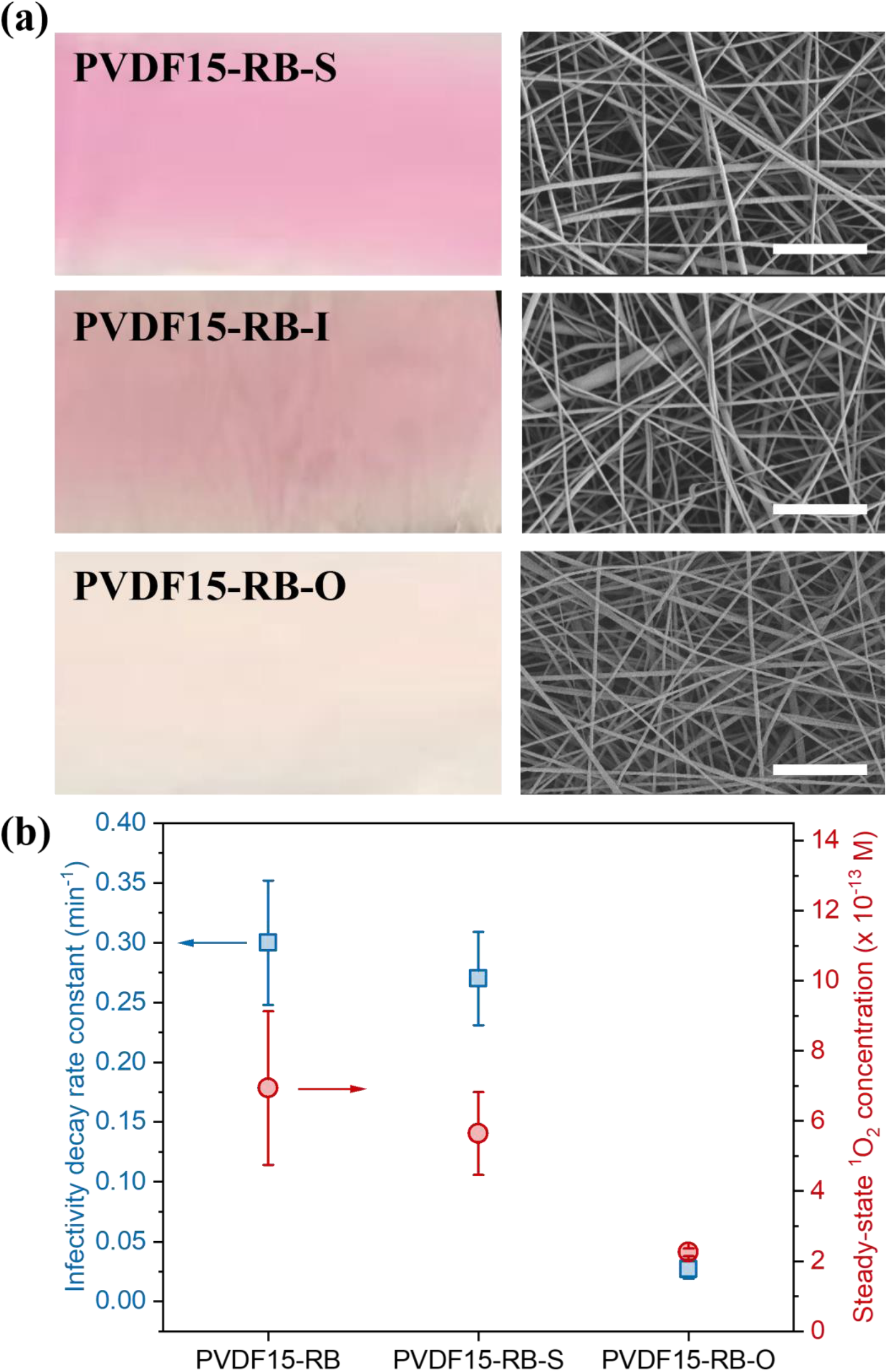
(a) Photos and SEM images of PVDF15-RB membranes after aging. Scale bars in the SEM images are 5 µm; (b) first-order decay rate constants of MHV-A59 infectivity in droplets and the corresponding ^1^O_2_ production on fresh and aged PVDF15-RB membranes. All tests of virus inactivation and ^1^O_2_ production were conducted under the irradiation of simulated reading light. Error bars for the infectivity decay rate constant represent the standard error of the negative slope of the linear regression of common logarithm of infectivity versus light exposure duration, and error bars for the steady-state ^1^O_2_ concentration represent the standard deviation of triplicate measurements.

Aged membranes were subjected for MHV-A59 aerosol filtration tests, and they all showed high filtration performance: the average filtration efficiency of PVDF15-RB-I, PVDF15-RB-S, and PVDF15-RB-O was 99.97, 99.96, and 99.95%, respectively. Aging under extended light exposure did not physically damage the membranes, and thus the aged membranes maintained excellent performance for removing MHV-A59 aerosols. However, aging did compromise membrane performance for coronavirus inactivation, particularly the significantly faded membrane after sunlight aging: PVDF15-RB-O inactivated MHV-A59 in droplets with *k*_*infectivity*_ only of 0.0273 ± 0.0079 min^-1^, which was 11.0 times lower than that of PVDF15-RB (0.301 ± 0.052 min^-1^, both tested under simulated reading light irradiation, *p* < 0.05, **Figure 3b**). In contrast, PVDF15-RB-S that did not fade after the aging by simulated indoor light kept its photoreactivity, and it inactivated MHV-A59 in droplets with *k*_*infectivity*_ of 0.268 ± 0.0386 min^-1^, which was comparable with that of PVDF15-RB (also tested under simulated reading light irradiation, *p* > 0.05). Fast coronavirus inactivation kinetics could be attributed to a higher [^1^O_2_]_ss_ produced by the membranes (**Figure 3b**). Since humans spend most of their time indoor and the risk of COVID-19 airborne transmission is much higher for indoor than outdoor activities, our results underscored that when used for face masks the photosensitized electrospun membranes could maintain their excellent performance of coronavirus aerosol filtration and coronavirus inactivation for a long duration under indoor light exposure.

## 4. Environmental Implication

Our study leverages nanotechnology to advance the design and fabrication of masks, respirators, and air filters, and it addresses the grand challenge of the airborne transmission of COVID-19. Photosensitized electrospun membranes showed excellent performance for capturing and killing coronavirus aerosols, i.e., they removed 99.2% of MHV-A59 aerosols, and inactivated 98.9% of MHV-A59 droplets only after 15 min of desk lamp irradiation. Electrospinning is an industrial viable and economically feasible technology for manufacturing new air filtration media at scale that could outperform current products on the market,^43,44^ such as cloth face masks and indoor air filters, for filtering out airborne pathogens. Miniaturized and portable electrospinning apparatuses further facilitate the wide deployment of the technique for individuals and small communities.^45^ Moreover, integration of dye photosensitizers as effective, low-cost, and biocompatible antimicrobial additives enables easy decontamination of air filtration media in and after use. Compared with many other disinfection strategies for reusing the masks and respirators in the COVID-19 pandemic, such as the treatment by heat, ultraviolet light irradiation, ozone, or hydrogen peroxide vapor,^46–48^ photosensitized electrospun air filters generate broad-spectrum biocides of ^1^O_2_ *in situ* under visible light exposure at the room temperature and pressure, and ^1^O_2_ could potentially damage multiple viral biomolecules and multiple steps in viral lifecycle to effectively inactivate coronaviruses. Photosensitized disinfection is less chemical and energy intensive, and it does not compromise the aerosol filtration efficiency after disinfection. In addition, color intensity and fading of the dye photosensitizers could serve as an indicator of the antimicrobial lifetime of the air filtration media.^49^

Future studies should focus on developing robust photoreactive air filters with extended lifetime to overcome the challenges of photobleaching of the dye-sensitized membranes, e.g., replacing the dyes with stable visible-light-responsive photocatalysts. In addition, further elucidating the mechanism of ROS damage to the coronaviruses will provide fundamental insights on designing advanced antimicrobial air filters.

## Acknowledgements

We acknowledge the NSF RAPID grants (CBET-2029411 and CBET-2029330) and the NSF grant (CBET-2028464) for supporting our research. We thank The George Washington University (GW) Nanofabrication and Imaging Center for SEM characterizations and GW BSL2+ lab facilities for the bioaerosol study. We thank Dominick J. Carluccio and Ryan Archer at the CH Technologies (USA), Inc. for characterizing aerosol size distribution.

## ASSOCIATED CONTENT

### Supporting Information

Experimental details; spectral irradiance, photon flux, and optical power density of the light sources; linear correlation of MHV-A59 infectivity with the viral load before infection; steady-state ^1^O_2_ concentrations produced by photosensitized electrospun air filters; aerosol size distribution; ORF5 gene damage of MHV-A59 aerosols in control experiments; infectivity of MHV-A59 droplets in control experiments.

