## Supporting Information for "Photosensitized Electrospun Nanofibrous Filters for Capturing and Killing Airborne Coronaviruses under Visible Light Irradiation"

Hongchen Shen,[1#] Zhe Zhou,[1#] Haihuan Wang,[1#] Mengyang Zhang,[1] Minghao Han,[2] Yun Shen,[2*] Danmeng Shuai[1*]

1 Department of Civil and Environmental Engineering, The George Washington University, Washington, DC 20052

2 Department of Chemical and Environmental Engineering, University of California, Riverside, Riverside, CA 92521

### Equal Contribution.

* Corresponding Authors:

Danmeng Shuai: Phone: 202-994-0506, Fax: 202-994-0127,, Website: http://materwatersus.weebly.com/

*Environmental Science & Technology*

8 Texts, 7 Figures, 1 Table, and 18 Pages

#### Materials and Methods

##### Test S1 Reagents and Materials

Polyvinylidene fluoride (PVDF) (Arkema KYNAR® 761), rose bengal (RB) (Sigma-Aldrich, dye content 95%), methylene blue hydrate (MB) (Sigma-Aldrich, 95+%), crystal violet (CV) (Sigma-Aldrich, 90+%), (-)-riboflavin (RF) (Sigma-Aldrich, 98+%), toluidine blue O (TBO) (Sigma-Aldrich, technical grade), *N*,*N*-dimethylformamide (DMF) (Sigma-Aldrich, 99+%), acetone (Sigma-Aldrich, 99.8+%), polypropylene (PP) fabrics (VWR® Basic Protection Face Mask and Amazon), Dulbecco's modified Eagle medium (DMEM, Corning®), fetal bovine essence (VWR), trypsin-ethylenediaminetetraacetic acid (trypsin-EDTA (0.25%), Gibco™), antibiotic antimycotic solution (Sigma-Aldrich, 100×), NaCl (VWR, ≥ 99.0%), and nuclease-free water (Fisher Scientific Inc.) were used as received, without further purification.

##### Text S2 Measurement of Dye Leaching

A whole sheet of PVDF15-RB after electrospinning was harvested and immersed in 150 mL of ultrapure water in a beaker. The system was next placed on an orbital shaker (120 rpm) for 3 h without light exposure. 1 mL of the aqueous solution was finally sampled at different time intervals for optical absorbance measurement by a UV-vis spectrophotometer (Hach DR6000) at 546 nm for RB.[1–3]

##### Text S3 Coronavirus Propagation

L-929 cells (ATCC CCL-1) were maintained with DMEM supplemented with 10% fetal bovine essence and 1% penicillin-streptomycin and incubated at 37 °C/5% $CO_2$ until confluence. Murine hepatitis virus A59 (MHV-A59) was inoculated in the confluent L-929 cells and incubated at 37 °C/5% $CO_2$ until a significant cytopathic effect was shown. After freeze-thaw for 3 times, all cell suspension was proceeded to 15 cycles of sonication in the ice-water mixture to liberate free MHV-A59 (each cycle consisted of 20 s of sonication on and 20 s of sonication off). Cell debris was removed by centrifugation (340 g, 10 min), and supernatant was aliquoted as the virus stock and stored at -80 °C until use.

**Text S4 Determination of Aerosol Filtration Efficiency**

A Blaustein atomizer (BLAM) was used to produce NaCl or MHV-A59 aerosols (air and liquid flow rate of ~1.5 L $min^{-1}$ and 15 mL $h^{-1}$, respectively), and single pass atomization was used to minimize virus damage. A portion of the aerosols with the flow rate of 0.5 standard liter per minute passed through a fitted membrane (11.0 $cm^2$) mounted on a filter holder and an impinger sequentially, and each filtration experiment (i.e., filter-on) was performed for 30 min. Aerosol size distribution was measured by Palas Promo 2000 (detection limit of 200 nm-10 μm) at the filter holder before filtration and reported as the number frequency. Aerosols that penetrated through the membrane were retained by the impinger prefilled with 4 mL of ultrapure water (for NaCl aerosols) or 4 mL of phosphate-buffered saline (PBS, for MHV-A59 aerosols). Control experiments (i.e., filter-off) were performed to quantify the aerosols collected in the impinger without an installed membrane.

**Text S5 RT-qPCR Quantification of Coronaviruses**

MHV-A59 collected in the impinger (**Text S4**) was first concentrated to ~100 µL by centrifugal ultrafiltration (Nanosep, 300 kDa, Pall Laboratory), and next proceeded for RNA extraction by a Zymo Quick-RNA Viral Kit (R1035). Both the centrifugal ultrafiltration membrane and the concentrate were used for RNA extraction to maximize the recovery of MHV-A59 in RNA extraction, and the recovery rate of 30.4 ± 1.6% was achieved.[4]

The TaqMan™ Fast Virus 1-Step Master Mix Kit (Thermo Fisher Scientific Inc., 4444432) was used for RT-qPCR. For each reaction of 20 µL, 5 µL of extracted RNA solution was mixed with 5 µL of Fast Virus Master Mix, 2 µL of forward primer (10 µM), 2 µL of reverse primer (10 µM), 1.25 µL of probe (10 µM), and 4.75 µL of RNA-free water. The primer was designed in the sequence of the ORF5 gene of the MHV-A59 structural protein M.[5,6] The hydrolysis probe consisted of fluorescein (FAM) at the 5' end and the Black Hole Quencher 1 dye (BHQ1) at the 3' end was utilized for quantification. Synthetic cDNA oligos (Integrated DNA Technologies, Inc.) in serial dilution ($10^3$-$10^{10}$ copies $mL^{-1}$) were used to plot the standard curve of the absolute gene copy numbers versus the threshold cycle (Ct) values. The RT-qPCR was run on the QuantStudio (Applied Biosystems) instrument with 10 min of reverse transcription at 52 ℃, 20 s of reverse transcription inactivation and denaturation initiation at 95 ℃, and 45 cycles of amplification at 95 °C for 15 s and 60 °C for 60 s. PCR amplification efficiency was determined as 86.7%-100.6%.[4] The virus stock and RNA-free water were used as the positive and negative control, respectively. Serial dilutions of extracted RNA from MHV-A59 did not indicate the presence of any inhibitors for qPCR.[4] All the RT-qPCR data were reported following MIQE guidelines.[4,7]

The sequences of primers were GGAACTTCTCGTTGGGCATTATACT and ACCACAAGATTATCATTTTCACAACATA; the sequence of the probe was 56-FAM/ACATGCTACGGCTCGTGTAACCGAACTGT/3BHQ_1, and the sequence of the cDNA standard was CTTAAGGAATGGAACTTCTCGTTGGGCATTATACTACTCTTTATTACTATCATACTAC AGTTCGGTTACACGAGCCGTAGCATGTTTATTTATGTTGTGAAAATGATAATCTTGT GGTTAATGTGGCC.[8]

**Text S6 Infectivity Determination of MHV-A59 Aerosols Captured on the Membrane**

The three-layer photosensitized electrospun membrane (i.e., sandwiched PVDF15-RB) immediately after filtering MHV-A59 aerosols was cut into six identical pieces and exposed to simulated reading light (**Figure S3**). One piece of the membrane was taken out at each time interval and vortexed in 3 mL of nuclease-free water for 3 min to release the virus. The 3 mL elution solution was then concentrated to ~100 µL by using centrifugal ultrafiltration (Nanosep, 300 kDa, Pall Laboratory) for the infectivity assay. A control experiment was conducted by dropping and drying a known amount of MHV-A59 solution on the membrane and subsequent RT-qPCR, and 54.6 ± 22.5% of the loaded virus was recovered.

50 µL of MHV-A59 after centrifugal ultrafiltration was introduced to 80-90% confluent L-929 cells in a 24-well plate and incubated at 37 °C/5% $CO_2$ for 2 h to allow efficient virus binding/entry but not significant replication. The inoculum was next removed and the cells were rinsed by 500

μL of pre-warmed DMEM (37 ℃) once. 500 μL of DMEM was then added into each well and the cells were incubated for another 24 h for RNA extraction and RT-qPCR quantification as indicated in **Text S5**.

**Text S7 Infectivity Determination of MHV-A59 Droplets**

Two-layer photosensitized electrospun membranes (i.e., PVDF15-RB) were first cut into several pieces and laid down at the bottom of a 24-well plate, and 200 μL of MHV-A59 solution (diluted from the virus stock by 100 times with nuclease-free water, ~$10^6$ gene copies $mL^{-1}$) was loaded on the membrane surface to form a thin layer of droplet (~ 1 mm) in each well. Next, the setup was exposed to simulated reading light and simulated indoor light, the virus solution was gently mixed on an orbital shaker (120 rpm), and 50 μL of the solution was sampled at each time intervals for the infectivity assay by ICC-RT-qPCR. After sampling, the membrane and the remaining virus solution were sacrificed.

**Text S8 Evaluation of Coronavirus' Lifecycle after Photosensitization**

For determining coronavirus binding, 50 μL of the virus solution was sampled at different time intervals, inoculated into 80-90% confluent L-929 cells together with 450 μL of DMEM in a 24-well plate, and incubated at 4 ℃ for 90 min. Specifically, the cells and DMEM were pre-cooled at 4 ℃ before incubation. Incubation at 4 ℃ allowed MHV-A59 binding to the cells but prevented its internalization and replication. Next, the cells were washed with 500 μL of ice-cold PBS once to remove unbound viruses, lysed for RNA extraction, and proceeded for RT-qPCR quantification of the ORF5 gene copy numbers of the viruses. For determining coronavirus internalization, the

L-929 cells after removing the unbound viruses were first introduced with 500 μL of pre-warmed DMEM (37 ℃) and incubated at 37 ℃ for another 90 min. Next, the cells were washed with trypsin-EDTA at 37 ℃ for 3 min to remove cell-bound but not internalized viruses, and the detached cells were centrifuged (340 g, 10 min), lysed for RNA extraction, and proceeded for RT-qPCR quantification of the ORF5 gene copy numbers of the viruses. Details of RNA extraction and RT-qPCR quantification are included in **Text S5**.

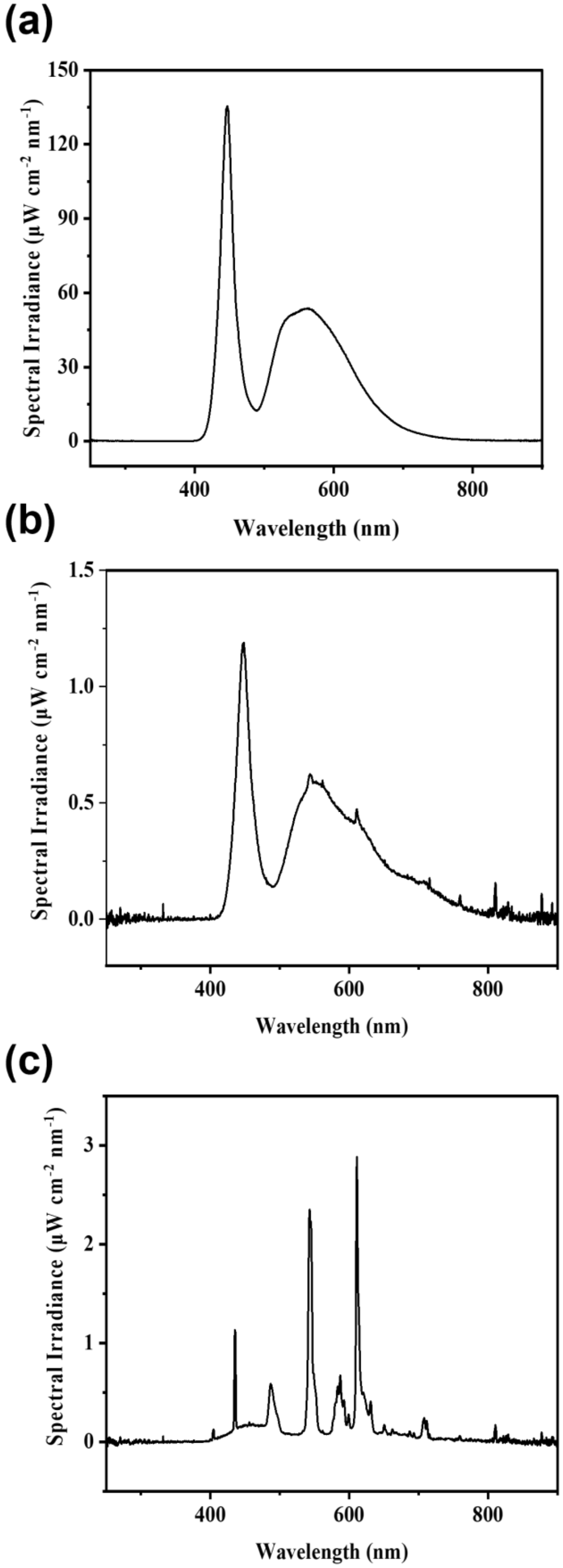

(a)
Spectral Irradiance (μW cm-2 nm-1)
Wavelength (nm)
(b)
Spectral Irradiance (μW cm-2 nm-1)
Wavelength (nm)
(c)
Spectral Irradiance (μW cm-2 nm-1)
Wavelength (nm)

**Figure S1.** The spectral irradiance of (a) simulated reading light (100% 7W white LED with a lamp-to-spectrometer detector distance of 15 cm); (b) simulated indoor light (3% 7W white LED with a lamp-to-spectrometer detector distance of 33 cm); and (c) indoor light (fluorescent light in the laboratory).

Outdoor aging for PVDF15-RB was conducted under the exposure of natural sunlight on The George Washington University campus, with sky conditions ranging from clear to partly cloudy (15:30, Nov 16$^{th}$, 2020 to 15:30, Nov 20$^{th}$, 2020).

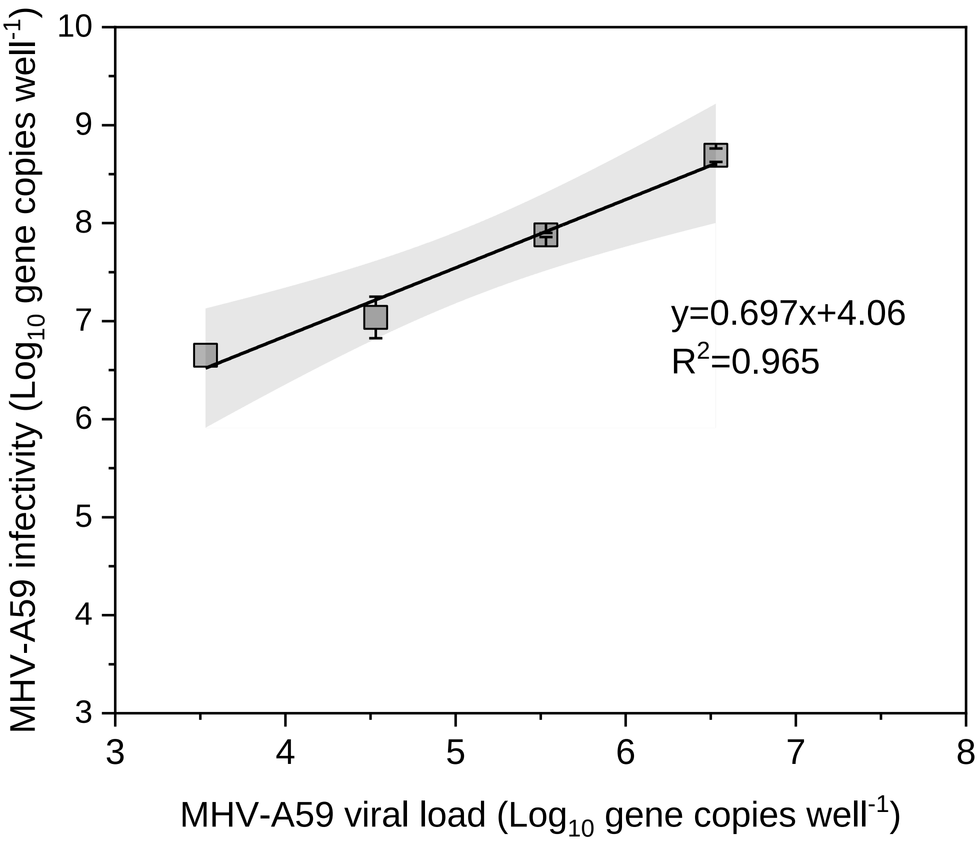


**Figure S2.** Linear correlation of MHV-A59 infectivity with the viral load before infection. The viral load was determined by RT-qPCR, and the infectivity was determined by ICC-RT-qPCR after 26 h of incubation with L-929 cells. Serial dilutions of the MHV-A59 virus stock were used to prepare different viral loads for cell infection. Special attention should be paid that the MHV-A59 gene input quantified by RT-qPCR did not indicate the concentration of infectious viruses. Error bars represent standard deviations, the black line stands for linear regression, and the light gray region highlights the 95% confidence band of the linear regression.

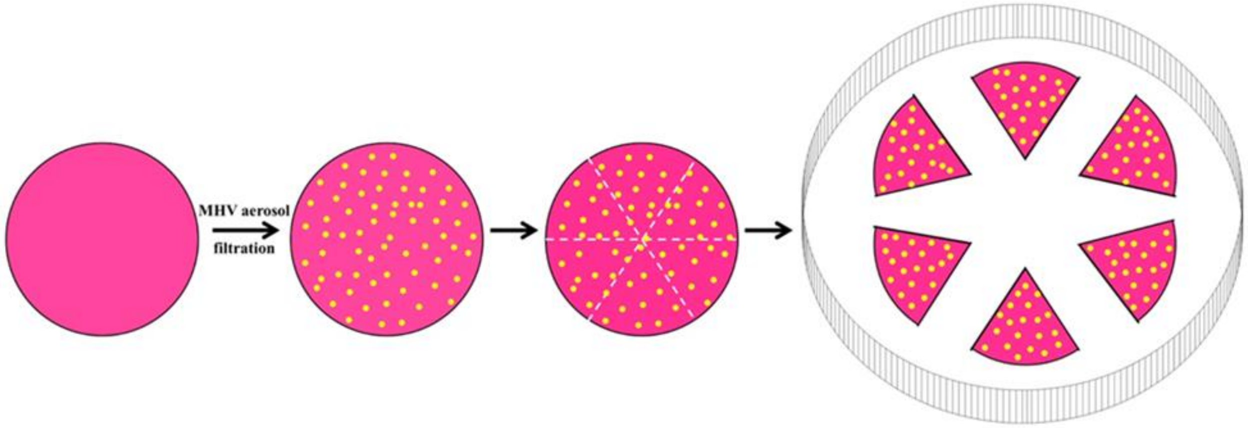


**Figure S3.** Schematic of PVDF15-RB and PVDF15 capturing MHV-A59 aerosols and subsequent evaluation of infectivity decay and gene damage after the irradiation of simulated reading light.

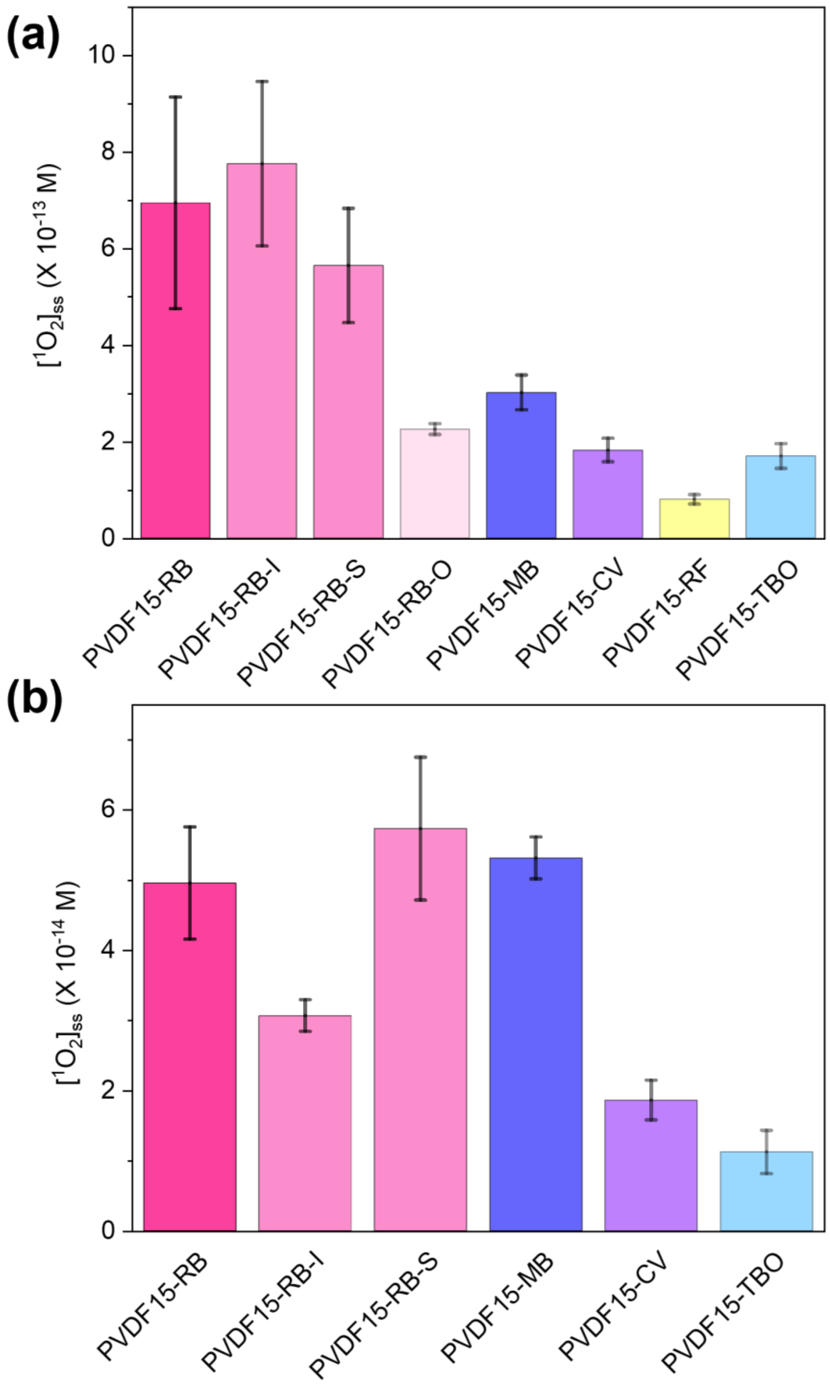


**Figure S4.** Steady-state $^1O_2$ concentrations produced by photosensitized electrospun membranes under the irradiation of (a) simulated reading light and (b) simulated indoor light. Error bars represent the standard deviation of triplicate measurements.

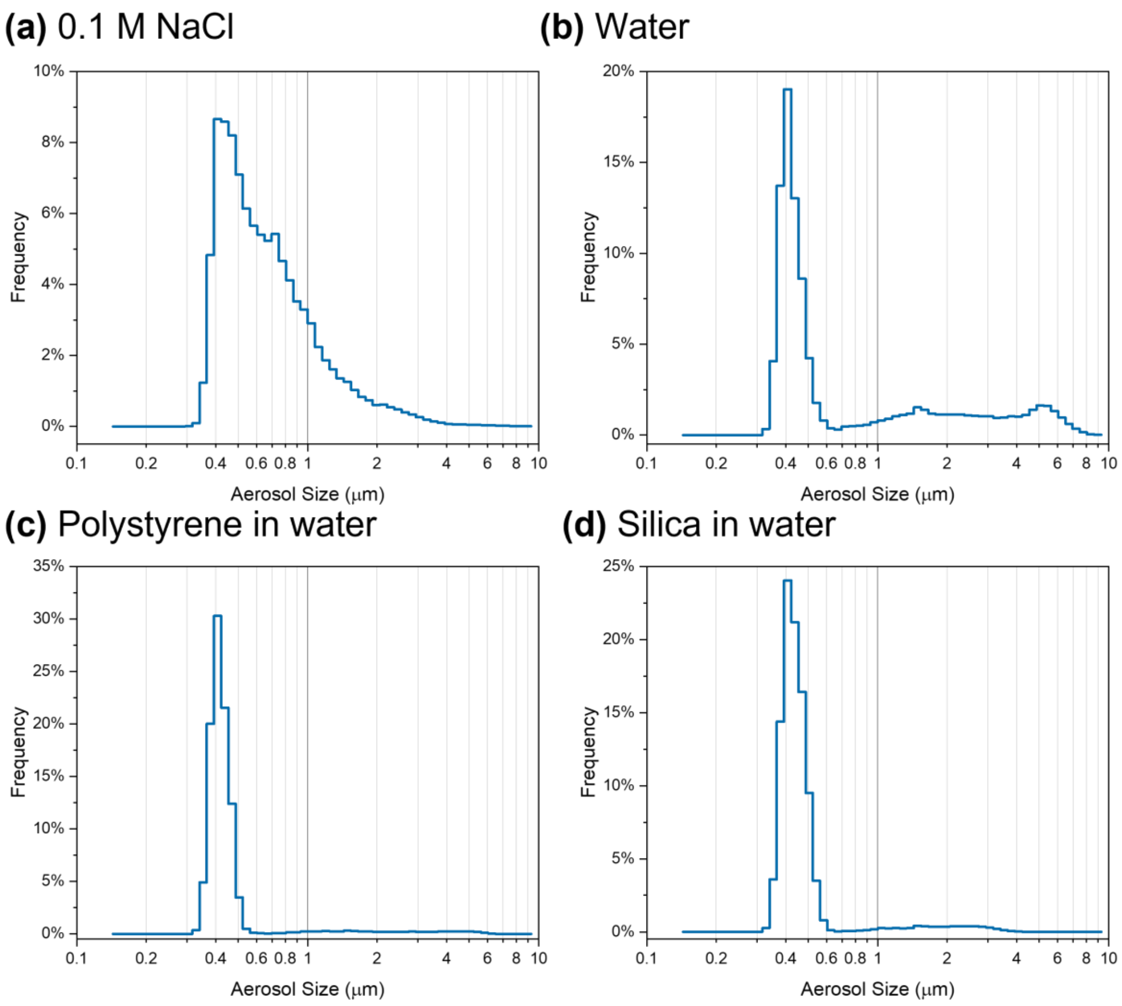


**Figure S5.** Size distribution of aerosols generated from (a) 0.1 M NaCl, (b) water, (c) 100 nm polystyrene particles in water, and (d) 100 nm silica particles in water. The number frequency of aerosols at different sizes was quantified by Palas Promo 2000 and was reported in the figures. Polystyrene and silica nanoparticles in water were used as surrogates for producing MHV-A59 aerosols, because they had a similar particle size with the virion size (100 nm of nanoparticles versus ~85 nm of MHV-A59), and their concentrations were similar (~$10^6$-$10^7$ particles $mL^{-1}$ versus ~$10^6$-$10^7$ gene copies $mL^{-1}$). The nanoparticles also had representative hydrophobicity/hydrophilicity that can simulate the virus. The most dominant aerosol size was 420-450 nm for all samples, which was within the range of the most penetrating aerosol size of

100-500 nm in air filtration. Palas Promo 2000 detected aerosols with a size of 200 nm-10 μm, and it was not able to quantify ultrafine aerosols below 100 nm that could be generated in our system. Data in (a) have been reported in our previous publication and it is shown here for reference.[4]

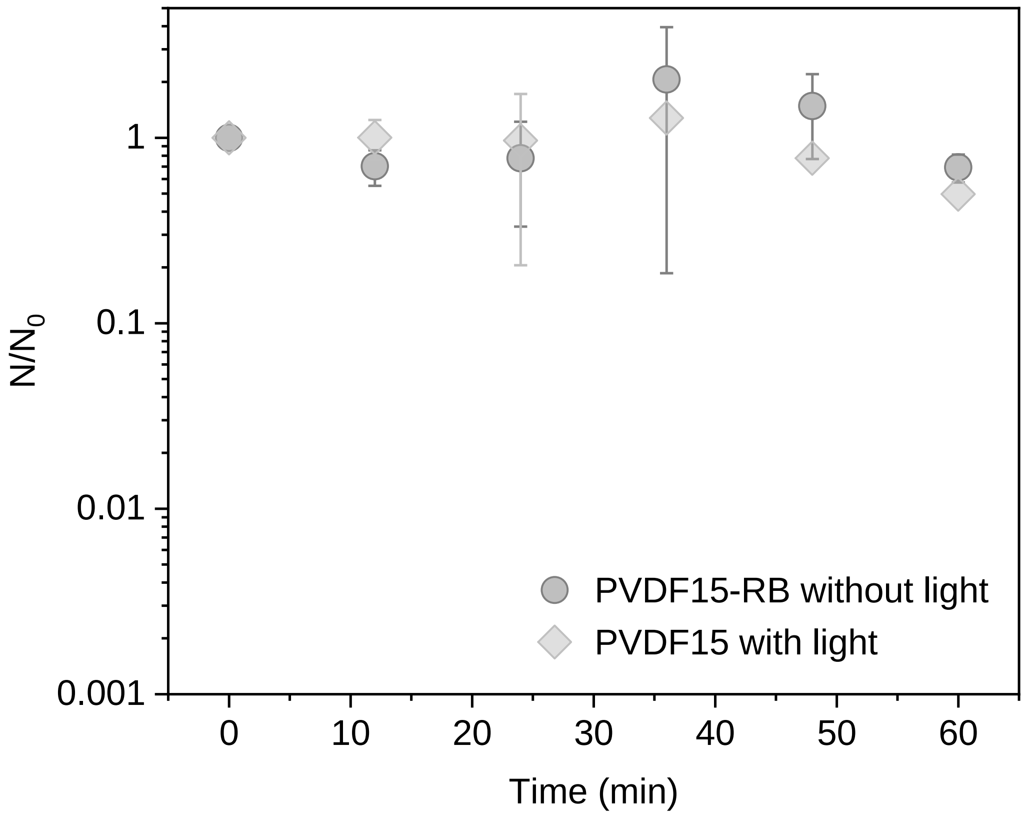


**Figure S6.** ORF5 gene damage kinetics of MHV-A59 aerosols captured on PVDF15-RB without light exposure (PVDF15-RB without light) as the dark control; and ORF5 gene damage kinetics of MHV-A59 aerosols captured on PVDF15 under the irradiation of simulated reading light (PVDF15 with light) as the light control. $N/N_0$ represents ORF5 gene copy numbers quantified by RT-qPCR at light exposure duration *t* to that at light exposure duration zero. Error bars represent standard deviations.

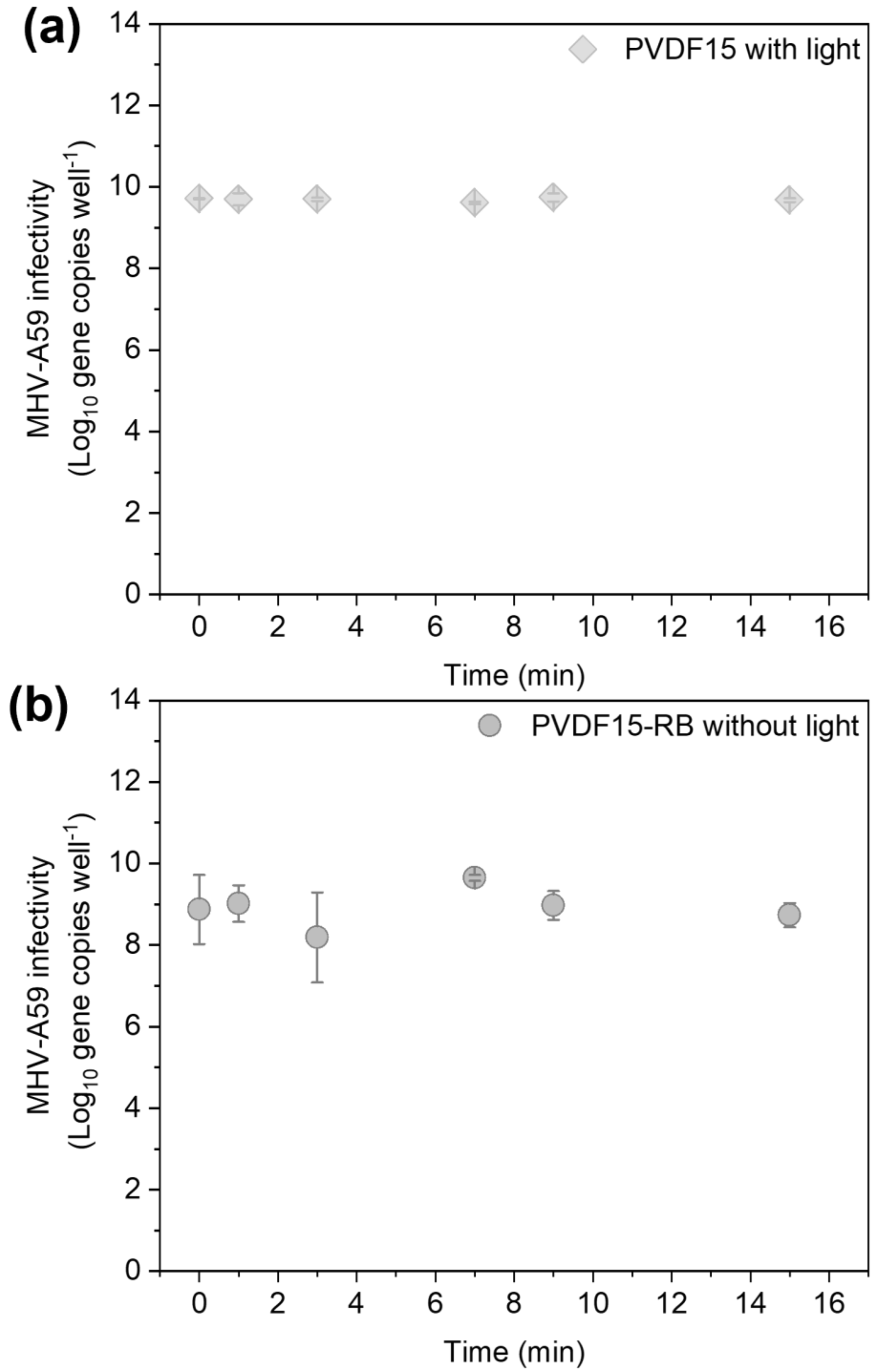


**Figure S7.** The infectivity of MHV-A59 droplets on (a) PVDF15 after exposure to simulated reading light and on (b) PVDF15-RB without light exposure. MHV-A59 infectivity was determined by ICC-RT-qPCR. Error bars represent standard deviations.

**Table S1.** Light source characterization

| Light source | Photon flux (μmol of photons $m^{-2}$ $s^{-1}$) | Optical power density (μWatt $cm^{-2}$) |
|---|---|---|
| Simulated reading light [a] | 590 | $1.32\times10^{4}$ |
| Simulated indoor light [b] | 4.94 | 106 |
| Indoor light [c] | 4.78 | 102 |

[a] The spectrometer detector was kept 15 cm away from the white LED lamp (7 W) for measuring the light intensity, and this lighting scenario represented strong reading lighting;

[b] The spectrometer detector was kept 33 cm away from the white LED lamp (7 W) after an optical filter (3 % light transmission, EDMUND OPTICS® 1.5 OD Absorptive ND filter), and this lighting scenario represented weak indoor lighting;

[c] Fluorescent light in the laboratory.
